## Supplementary Information for "Structural basis of substrate recognition and membrane association by the bacterial lysyl-phosphatidylglycerol hydrolase AcvB"

Supplementary items:

Supplementary Figures 1–3

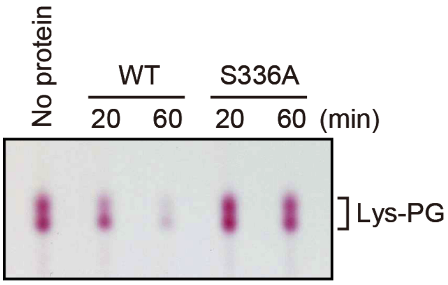


**Supplementary Fig. 1: Purified AcvB(25–456) exhibits Lys-PG hydrolase activity.**
Purified wild-type (WT) and S336A-mutant AcvB(25–456) were incubated with Lys-PG at 37 °C for the indicated times. Lipids were extracted, separated through TLC, and visualized using ninhydrin staining.

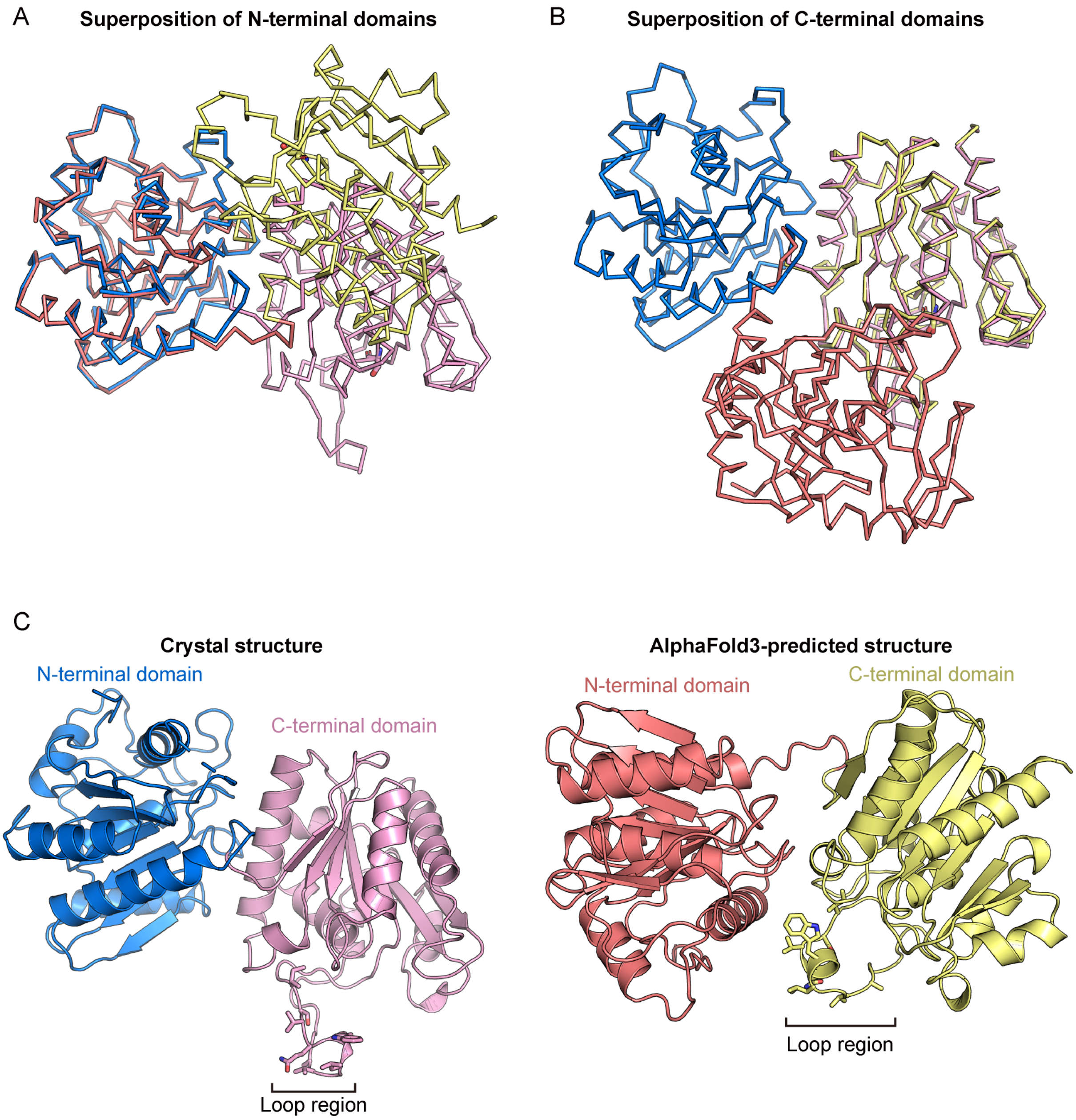


**Supplementary Fig. 2: Comparison of the crystal and AlphaFold3-predicted structures of AcvB(25–456).**

(A) Superposition of the N-terminal domains of AcvB(25–456) in the crystal structure and the model predicted by AlphaFold3. The N-terminal and C-terminal domains of the crystal structure are colored blue and pink, respectively, whereas those of the AlphaFold3-predicted model are colored red and yellow, respectively. (B) Superposition of the C-terminal domains of AcvB(25–456) in the crystal structure and AlphaFold3-predicted model. (C) Crystal structure (left) and AlphaFold3-predicted structure (right). The hydrophobic protruding loop regions are indicated.

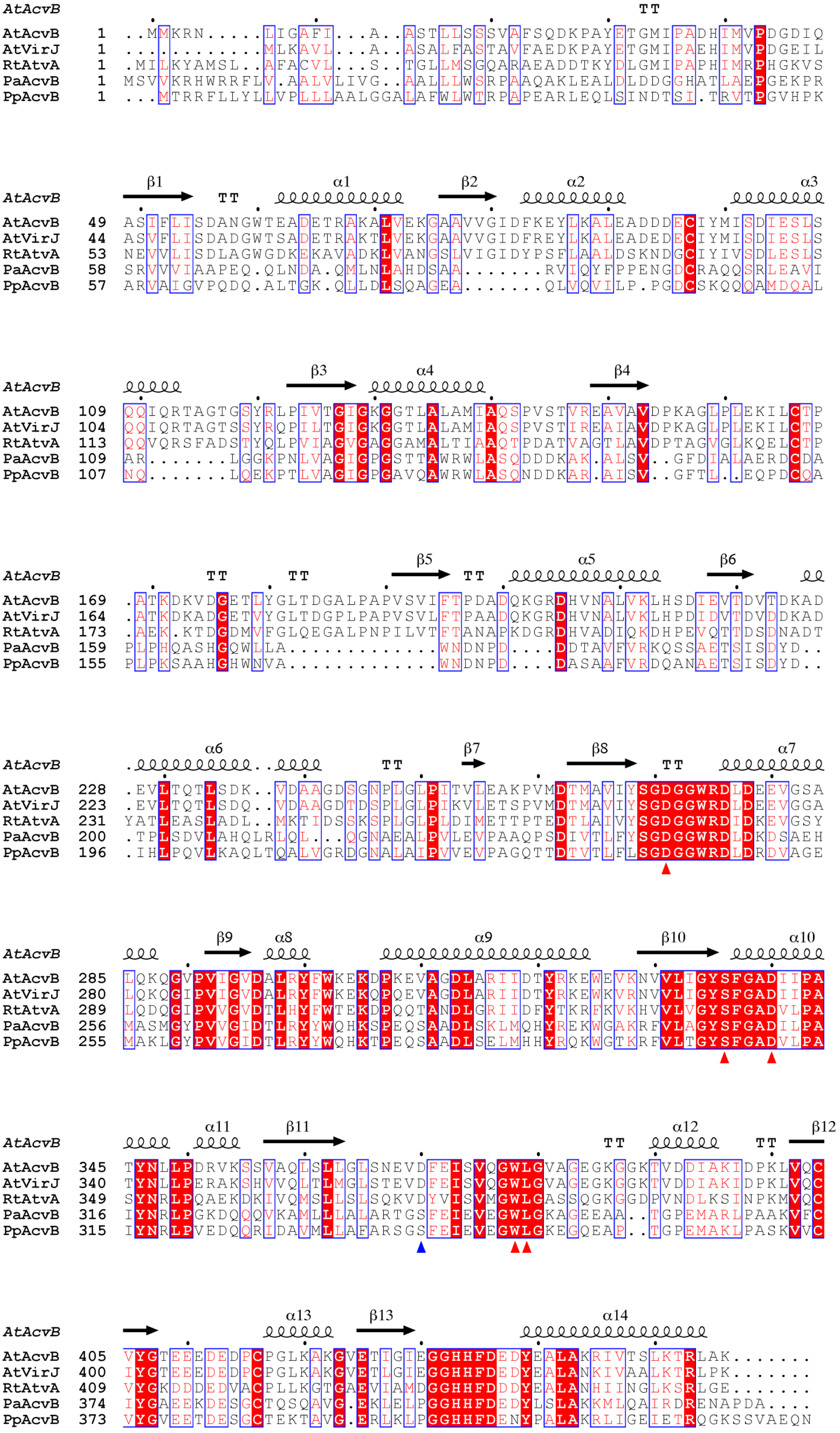
AtAcvB TT
AtAcvB 1 ..MMKRN.....LIGAFI...A..ASTLLSSSVAFSQDKPAYETGMIPADHIMVPDGDIQ
AtVirJ 1 ...........MLKAVL...A..ASALFASTAVFAEDKPAYETGMIPAEHIMVPDGEIL
RtAtvA 1 .MILKYAMSL..AFACVL...S..TGLLMSGQARAEADDTKYDLGMIPAPHIMRPHGKVS
PaAcvB 1 MSVVKRHWRRFLV.AALVLIVG..AALLLWSRPAAQAKLEALDLDDGGHATLAEPGEKPR
PpAcvB 1 ...MTRRFLLYLLVPLLLAALGGALAFWLWTRPAPEARLEQLSINDTSI.TRVTPGVHPK
β1 TT α1 β2 α2 α3
AtAcvB 49 ASIFLISDANGWTEADETRAKALVEKGAAVVGIDFKEYLKALEADDDECIYMISDIESLS
AtVirJ 44 ASVFLISDADGWTSADETRAKTLVEKGAAVVGIDFREYLKALEADEDECIYMISDIESLS
RtAtvA 53 NEVVLISDLAGWGDKEKAVADKLVANGSLVIGIDYPSFLAALDSKNDGCIYIVSDLESLS
PaAcvB 58 SRVVVIAAPEQ.QLNDA.QMLNLAHDSAA.......RVIQYFPPENGDCRAQQSRLEAVI
PpAcvB 57 ARVAIGVPQDQ.ALTGK.QLLDLSQAGEA.......QLVQVILP.PGDCSKQQQAMDQAL
β3 α4 β4
AtAcvB 109 QQIQRTAGTGSYRLPIVTGIGKGGTLALAMIAQSPVSTVREAVAVDPKAGLPLEKILCTP
AtVirJ 104 QQIQRTAGTSSYRQPILTGIGKGGTLALAMIAQSPVSTIREAIAVDPKAGLPLEKILCTP
RtAtvA 113 QQVQRSFADSTYQLPVIAGVGAGGAMALTIAAQTPDATVAGTLAVDPTAGVGLKQELCTP
PaAcvB 109 AR.......LGGKPNLVAGIGPGSTTAWRWLASQDDDKAK.ALSV..GFDIALAERDCDA
PpAcvB 107 NQ.......LQEKPTLVAGIGPGAVQAWRWLASQNDDKAR.AISV..GFTL..EQPDCQA
TT TT β5 TT α5 β6
AtAcvB 169 .ATKDKVDGETLYGLTDGALPAPVSVIFTPDADQKGRDHVNALVKLHSDIEVTDVTDKAD
AtVirJ 164 .ATKDKADGETVYGLTDGPLPAPVSVLFTPAADQKGRDHVNALVKLHPDIDVTDVDDKAD
RtAtvA 173 .AEK.KTDGDMVFGLQEGALPNPILVTFTANAPKDGRDHVADIQKDHPEVQTTDSDNADT
PaAcvB 159 PLPHQASHGQWLLA.............WNDNPD....DDTAVFVRKQSSAETSISDYD..
PpAcvB 155 PLPKSAAHGHWNVA.............WNDNPD....DASAAFVRDQANAETSISDYD..
α6 TT β7 β8 TT α7
AtAcvB 228 .EVLTQTLSDK..VDAAGDSGNPLGLPITVLEAKPVMDTMAVIYSGDGGWRDLDEEVGSA
AtVirJ 223 .EVLTQTLSDQ..VDAAGDTDSPLGLPIKVLETSPVMDTMAVIYSGDGGWRDLDEEVGGA
RtAtvA 231 YATLEASLADL..MKTIDSSKSPLGLPLDIMETTPTEDTLAIVYSGDGGWRDIDKEVGSY
PaAcvB 200 .TPLSDVLAHQLRLQL...QGNAEALPVLEVPAAQPSDIVTLFYSGDGGWRDLDKDSAEH
PpAcvB 196 .IHLPQVLKAQLTQALVGRDGNALAIPVVEVPAGQTTDTVTLFLSGDGGWRDLDRDVAGE
β9 α8 α9 β10 α10
AtAcvB 285 LQKQGVPVIGVDALRYFWKEKDPKEVAGDLARIIDTYRKEWEVKNVVLIGYSFGADIIPA
AtVirJ 280 LQKQGIPVIGVDALRYFWKEKQPQEVAGDLARIIDTYRKEWKVRNVVLIGYSFGADIIPA
RtAtvA 289 LQDQGIPVVGVDTLHYFWTEKDPQQTANDLGRIIDFYTKRFKVKHVVLVGYSFGADVLPA
PaAcvB 256 MASMGYPVVGIDTLRYYWQHKSPEQSAADLSKLMQHYREKWGAKRFVLAGYSFGADILPA
PpAcvB 255 MAKLGYPVVGIDTLRYYWQHKTPEQSAADLSELMHHYRQKWGTKRFVLTGYSFGADVLPA
α11 β11 TT α12 TT β12
AtAcvB 345 TYNLLPDRVKSSVAQLSLLGLSNEVDFEISVQGWLGVAGEGKGGKTVDDIAKIDPKLVQC
AtVirJ 340 TYNLLPERAKSHVVQLTLMGLSTEVDFEISVQGWLGVAGEGKGGKTVDDIAKIDPKLVQC
RtAtvA 349 SYNRLPQAEKDKIVQMSLLSLSQKVDYVISVMGWLGASSQGKGGDPVNDLKSINPKMVQC
PaAcvB 316 IYNRLPGKDQQQVKAMLLLALARTGSFEIEVEGWLGKAGEEAA..TGPEMARLPAAKVFC
PpAcvB 315 IYNRLPVEDQQRIDAVMLLAFARSGSFEIEVEGWLGKEGQEAP..TGPEMAKLPASKVVC
α13 β13 α14
AtAcvB 405 VYGTEEEDEDPCPGLKAKGVETIGIEGGHHFDEDYEALAKRIVTSLKTRLAK.......
AtVirJ 400 IYGTEEEDEDPCPGLKAKGVETLGIEGGHHFDEDYEALANKIVAALKTRLPK.......
RtAtvA 409 VYGKDDDEDVACPLLKGTGAEVIAMDGGHHFDDDYEALANHIINGLKSRLGE.......
PaAcvB 374 IYGAEEKDESGCTQSQAVG.EKLELPGGHHFDEDYLSLAKKMLQAIRDRENAPDA....
PpAcvB 373 VYGVEETDESGCTEKTAVG.ERLKLPGGHHFDENYPALAKRLIGEIETRQGKSSVAEQN

**Supplementary Fig. 3: Sequence alignment of AcvB homologs.**

Multiple-sequence alignment of *Agrobacterium tumefaciens* AcvB (AtAcvB) and its homologs: *A. tumefaciens* VirJ (AtVirJ), *Rhizobium tropici* AtvA (RtAtvA), PA0919 from *Pseudomonas aeruginosa* (PaAcvB), and PP_1201 from *Pseudomonas putida* (PpAcvB). Identical residues are shaded red, similar residues are colored red, and similar residues across groups are boxed in blue. The secondary structure of AcvB is shown above the sequences. Conserved residues Asp271, Ser336, Asp340, Trp378, and Leu379 in AcvB are indicated by red arrowheads, and Asp370 is indicated by a blue arrowhead.
